# Short single-stranded DNA oligonucleotides enable intracellular transcription of functional RNAs in mammalian cells

**DOI:** 10.64898/2026.05.30.729002

**Authors:** Ahmed Mahas, Ramiro M. Perrotta, Roman Teo Oliynyk, Raphael Ferreira, George M. Church

## Abstract

Programmable intracellular production of short functional RNAs underlies diverse applications ranging from gene regulation to genome engineering, but is often constrained by cloning workflows, RNA synthesis, or viral-vector delivery. Here, we introduce a single-stranded DNA (ssDNA)-driven intracellular transcription strategy in which short ssDNA templates function as transcriptional substrates when paired with orthogonal bacteriophage RNA polymerases (RNAPs). By embedding phage promoters within short hairpin structures to create a locally double-stranded recognition site, we enable robust RNAP-dependent transcription from ssDNA in mammalian cells. We demonstrate that ssDNA templates can drive production of functional CRISPR guides, supporting adenine base editing at reporter and endogenous loci and enabling CRISPR-based transcriptional activation across distinct human cell lines. These results establish ssDNA templates as compact, synthetically accessible inputs for RNA production in mammalian cells, with broad utility in synthetic biology and genome engineering.

## Introduction

Efficient intracellular production of short functional RNAs is essential for many mammalian cell engineering applications, ranging from genome editing to programmable gene regulation. Common delivery modalities, including plasmid DNA, in vitro–transcribed RNA, synthetic RNA oligonucleotides, and viral vectors, are effective but impose practical constraints such as cloning requirements, scalability limitations, or challenges in RNA stability and delivery. These constraints highlight the need for a simple, modular way to generate short programmable RNAs intracellularly from minimal, cloning-free, and non-viral templates.

ssDNA is an attractive candidate for such an approach because it can be synthesized rapidly and inexpensively with high sequence fidelity, is easily multiplexed, and is chemically tunable. However, short ssDNA templates have not been established as transcriptional substrates in mammalian cells. Bacteriophage RNA polymerases (RNAPs) provide a potential solution because they recognize short promoter sequences (1) and operate orthogonally to mammalian transcriptional machinery.

Here, we show that engineered ssDNA templates can function as transcriptionally competent expression substrates in mammalian cells when coupled to RNAPs. By embedding phage promoters within short hairpin structures to provide a locally double-stranded recognition site, we enable robust phage RNAP–dependent transcription from ssDNA, including the production of functional gRNAs that support genome editing and programmable transcriptional activation.

## Results

To test whether ssDNA can function as a transcriptional expression template in mammalian cells, we designed minimal ssDNA constructs encoding a bacteriophage promoter followed by a CRISPR guide sequence to provide a simple, activity-based readout of RNA output (Fig. 1A). The promoter region was configured as a short hairpin structure to provide a locally double-stranded substrate for bacteriophage RNAP recognition, while the downstream sequence remained single-stranded (2).

**Figure 1.**
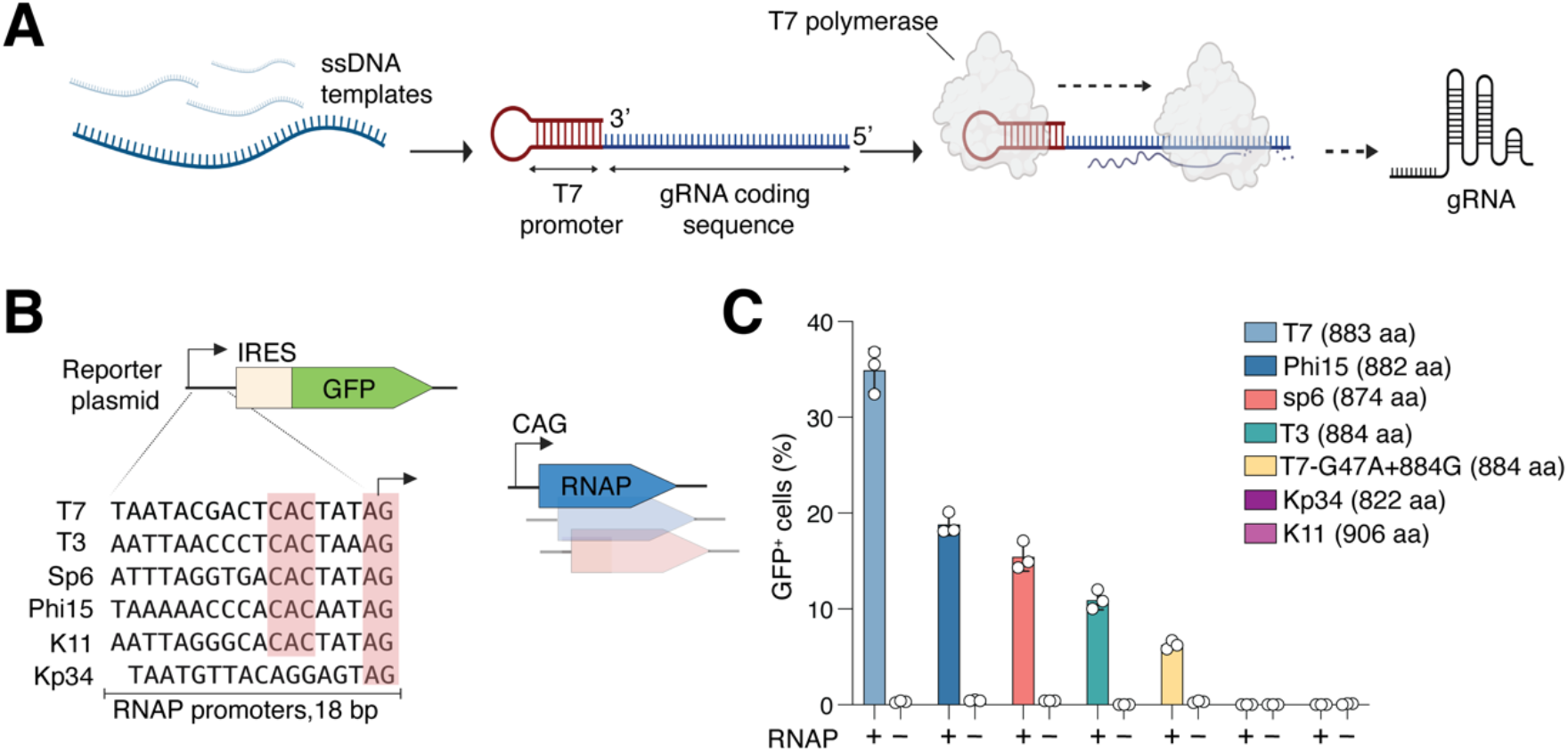
Distinct bacteriophage RNA polymerases drive different levels of gene expression in mammalian cells. **(A)** Schematic illustrating the ssDNA-based transcription strategy. ssDNA templates encoding a hairpin-configured bacteriophage promoter upstream of a short functional RNA sequence are delivered into mammalian cells together with bacteriophage RNA polymerase (e.g., T7 polymerase). The promoter hairpin provides a locally double-stranded recognition site, enabling RNAP-dependent transcription of functional RNA (e.g., gRNAs) from ssDNA templates. **(B)** Sequence alignment of bacteriophage RNAP promoter variants used in this study, including T7, T3, SP6, Phi15, K11, and Kp34. Conserved core promoter regions critical for RNAP recognition are highlighted. IRES, internal ribosomal entry site. **(C)** Quantification of GFP expression from plasmid reporters using different bacteriophage RNAPs. GFP expression is observed only in the presence of the corresponding RNAP, demonstrating strict polymerase dependence and orthogonality. Bars represent mean ± s.d. from independent biological replicates, *n* = 3.

Before testing ssDNA templates directly, and to establish baseline activity and enable direct comparison across polymerase–promoter pairs, we first evaluated the functionality of different RNAPs in human cells to drive transcription from plasmid templates using a green fluorescent protein (GFP) reporter assay. We examined promoter sequences recognized by multiple RNAPs, including T7 (3) (WT and engineered T7-G47A+884G (4)), T3 (5), SP6 (6), Phi15 (7), K11 (8), and KP34 (9) (Fig. 1B). Among the polymerases tested, T7 RNAP produced the highest transcriptional output, while Phi15, SP6, and T3 RNAPs supported lower but detectable levels of expression. No GFP signal was observed in the absence of RNAP, confirming the orthogonality of the system (Fig. 1C).

We next tested whether these ssDNA templates support RNAP-dependent intracellular transcription that yields functional RNA output. To address this, we coupled ssDNA-driven gRNA expression to a CRISPR adenine base editor (ABE (10)) reporter in which editing activity restores mCherry expression (Fig. 2A). Delivery of ssDNA templates encoding a targeting gRNA produced clear reporter activation above the non-targeting control, indicating functional gRNA output from ssDNA in mammalian cells (Fig. 2B). Notably, the optimized configuration increased the fraction of mCherry+cells from 6.7% (HL, no PS) to 17.4% (HL+PS). Importantly, T7–NLS outperformed T7+NLS by ~1.6×, indicating that cytoplasmic-accessible polymerase enhances ssDNA-driven output (Fig. 2B).

**Figure 2.**
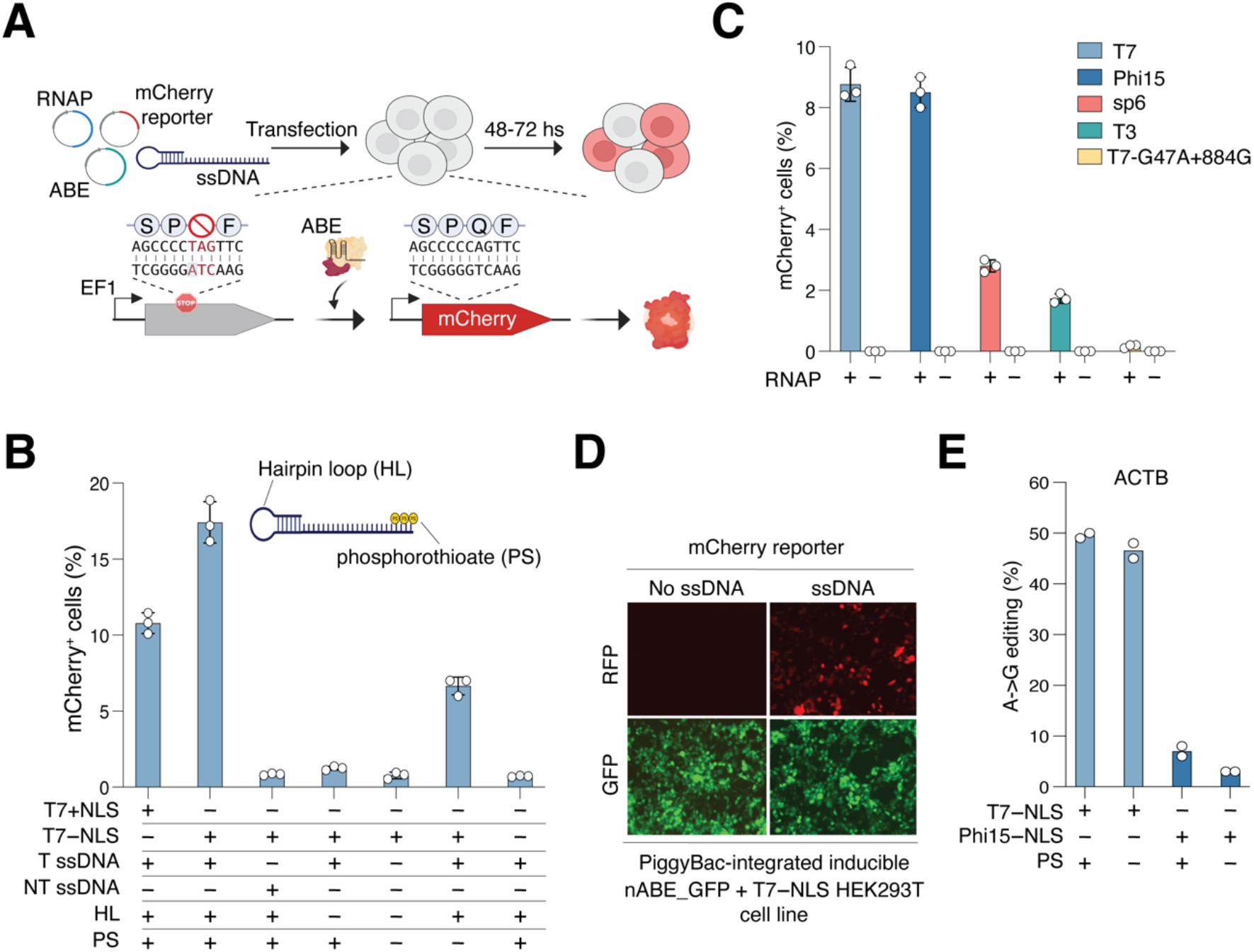
Single-stranded DNA templates enable bacteriophage RNA polymerase– dependent transcription and genome editing in mammalian cells. **(A)** Experimental workflow for functional assessment of ssDNA-driven gRNA expression using a CRISPR adenine base editing reporter system. ssDNA templates encoding gRNAs are co-delivered with RNAPs and ABE to edit an engineered stop codon within the mCherry-coding sequence, leading to reporter activation upon successful gRNA expression and editing. **(B)** Evaluation of ssDNA design features affecting functional output. Incorporation of a promoter hairpin loop (HL) and terminal phosphorothioate (PS) modifications enhances ssDNA-driven editing efficiency, defining design principles for robust intracellular transcription. Bars represent mean ± s.d. from independent biological replicates, *n* = 3. NLS: nuclear localization signal. T ssDNA: ssDNA encoding targeting gRNA. NT ssDNA: ssDNA encoding non-targeting gRNA. **(C)** Quantification of reporter activation following ssDNA-driven gRNA expression using different RNAPs. Editing activity mirrors transcriptional output and is strictly RNAP-dependent. Bars represent mean ± s.d. from independent biological replicates, *n* = 3. **(D)** Representative fluorescence microscopy images showing reporter activation in cells engineered to express T7 RNAPs and ABE transfected with or without ssDNA templates. Red fluorescence indicates editing-dependent reporter activation, while green fluorescence marks the dox-inducible expression of ABE. **(E)** Endogenous genome editing at the ACTB locus mediated by ssDNA-driven gRNA expression. A-to-G editing efficiencies are shown for T7 and Phi15 RNAPs, with and without PS stabilization. Bars represent mean ± s.d. from independent biological replicates, *n* = 2.

We next evaluated whether the ssDNA strategy extends to additional phage RNAPs beyond T7. While T7 polymerase was most active, Phi15, SP6, and T3 RNAPs supported measurable RNAP-dependent reporter activation, indicating that ssDNA templates can function across distinct phage promoter–polymerase pairs (Fig. 2C).

To assess whether ssDNA-driven gRNA expression could be integrated into more scalable workflows, we generated HEK293T cell lines with PiggyBac-integrated, doxycycline-inducible T7 RNAP and ABE. After induction, transfection with mCherry reporter and gRNA-encoding ssDNA templates resulted in editing-dependent reporter activation, demonstrating that ssDNA-derived gRNAs function in cell lines in which the core components are stably encoded (Fig. 2D).

Next, we assessed whether ssDNA-mediated transcription could support genome editing at an endogenous genomic locus. Using ssDNA templates encoding ACTB-targeting gRNAs, we observed efficient adenine-to-guanine editing in mammalian cells in an RNAP-dependent manner (Fig. 2E). Editing efficiencies were highest for T7 RNA polymerase and were enhanced by PS stabilization, consistent with reporter-based results.

To extend ssDNA-driven transcription beyond genome editing, we next tested whether it can drive CRISPR-based transcriptional activation (CRISPRa). We co-delivered T7 RNAP, a dCas12-based transcriptional activator (dCas12–mVPR)(11), and ssDNA templates encoding a CRISPR RNA (crRNA) designed to recruit dCas12–mVPR to a TRE3G reporter containing seven TetO repeats upstream of GFP (Fig. 3A). In HEK293T cells, delivery of ssDNA templates encoding targeting crRNAs resulted in robust GFP activation (~40–45% GFP^+^ cells), whereas a non-targeting control yielded background-level GFP signal (Fig. 3B). To assess generalizability across cell types, we performed the same ssDNA-driven CRISPRa assay in HeLa cells and observed reproducible GFP induction above the non-targeting control (~5–6% GFP^+^ cells) (Fig. 3C).

**Figure 3.**
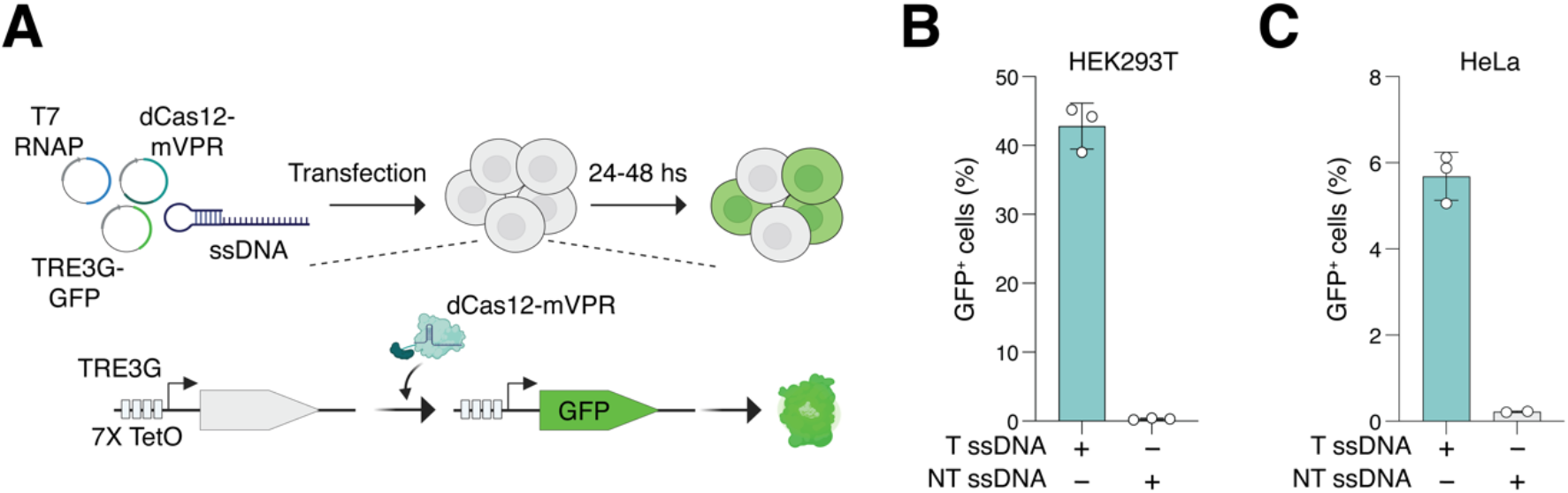
ssDNA-driven crRNA expression enables CRISPR-based transcriptional activation in mammalian cells. **(A)** Schematic of the ssDNA-driven CRISPR activation (CRISPRa) assay. ssDNA templates encoding a targeting crRNA are co-delivered with T7 RNA polymerase (T7 RNAP) and a dCas12–mVPR transcriptional activator to recruit the activator to a TRE3G promoter containing seven TetO repeats upstream of a GFP reporter, resulting in GFP induction upon successful crRNA expression and targeting. **(B)** Quantification of GFP activation following ssDNA-driven CRISPRa in HEK293T and **(C)** HeLa cells. Targeting crRNA templates (T ssDNA) drive robust GFP induction relative to the non-targeting control crRNA (NT ssDNA), demonstrating functional crRNA transcription from ssDNA templates across distinct mammalian cell types. Bars represent mean ± s.d. from independent biological replicates, *n* = 3 (*n* = 2 in NT ssDNA in Fig. 3C).

## Discussion

Together, these results establish short ssDNA templates as transcriptionally competent expression substrates in mammalian cells when paired with orthogonal bacteriophage RNAPs. By converting compact synthetic DNA oligonucleotides into functional short RNAs inside cells, this approach provides a simple alternative to plasmid- or RNA-based delivery that is readily programmable through sequence design and template chemistry.

Notably, ssDNA is already widely used in genome engineering workflows as a scalable and chemically tunable nucleic acid format, with established synthesis pipelines, quality control, and straightforward multiplexing (12). Leveraging these practical advantages for intracellular RNA expression could enable rapid design–test cycles and facilitate pooled or combinatorial perturbation strategies without requiring molecular cloning or in vitro transcription. Because guide templates can be synthesized as standardized oligonucleotide libraries, this framework is naturally compatible with multiplexed and automated workflows where many guide sequences or conditions must be evaluated in parallel.

Beyond genome editing, we show that ssDNA-driven guide expression supports programmable transcriptional activation across distinct human cell lines, highlighting the broader utility of this expression modality for CRISPR regulation. Future work may extend ssDNA-templated transcription to additional polymerases, including endogenous mammalian polymerases. In parallel, this framework could enable sophisticated control of guide expression timing and localization for synthetic biology applications.

## Methods

### Single-stranded DNA designs and generation

All ssDNA oligos were synthesized and obtained from Integrated DNA Technologies (IDT) as ultramers with or without 5’ end phosphorothioate modifications. The T7 promoter– containing ssDNA template used for the ABE/mCherry reporter experiments (5′→3′) was: (c*t*agcaccgactcggtgccactttttcaagttgataacggactagccttattttaacttgctatttctagctctaaaac<u>gggac atcctgagcccctag</u>CCTATAGTGAGTCGTATTAatttcgcgtgtttgtcgcgaaatTAATACGACTCAC TATAGG).

The ACTB-targeting ssDNA template (5′→3′) was: (c*t*agcaccgactcggtgccactttttcaagttgataacggactagccttattttaacttgctatttctagctctaaaact<u>ggtga gctgcgagaatag</u>CCTATAGTGAGTCGTATTAatttcgcgtgtttgtcgcgaaatTAATACGACTCACT ATAGG). For CRISPRa experiments, the ssDNA template encoding a dCas12 crRNA targeting TetO (5′→3′) was: (<u>c*g*ttctctatcactgatagggag</u>atctacacttagtagaaattaattatCCCTATAGTGAGTCGTATTAatttcg cgtgtttgtcgcgaaatTAATACGACTCACTATAGGG) and the non-targeting (NT) control ssDNA (5′→3′) was: (<u>cgtcatctctgaccctcttttag</u>atctacacttagtagaaattaattatCCCTATAGTGAGTCGTATTAatttcgcgt gtttgtcgcgaaatTAATACGACTCACTATAGGG). Asterisks (*) indicate phosphorothioate (PS) backbone linkages and the underlined region corresponds to the guide/crRNA spacer sequence. Uppercase bases indicate the T7 promoter sequence. For experiments using alternative phage RNA polymerases, the T7 promoter segment was replaced with the corresponding promoter variants listed in Fig. 1B.

### Plasmids

All plasmids were generated using standard molecular cloning methods, including Gibson assembly and restriction–ligation cloning. The different RNAPs coding sequences were human codon optimized and obtained as synthetic gBlocks (IDT). The gBlocks were then assembled under the control of the CAG promoter into the backbone of Addgene plasmid #65974 (digested with EcoRI and NheI) using the NEB HiFi DNA Assembly Master Mix.

To generate GFP reporter constructs driven by different RNAPs promoters, we used the T7-driven GFP reporter plasmid (Addgene #138586) as either the T7 promoter construct or as the cloning backbone for alternative promoter variants. For non-T7 promoters, synthetic promoter sequences were introduced by replacing the T7 promoter region in Addgene #138586 with double-stranded oligonucleotides (IDT). Complementary oligos containing the desired promoter sequences were phosphorylated and annealed using T4 Polynucleotide Kinase (NEB), and the resulting duplexes were ligated into XmaI- and NheI-digested #138586 backbone using T4 DNA Ligase (NEB).

To generate the mCherry reporter plasmid, a premature stop codon was introduced by mutating the cytosine (C) in the glutamine codon at position 69 to thymine (T). The modified mCherry coding sequence was synthesized as a gBlock (IDT) and assembled under the control of an EF1α promoter into the backbone of Addgene plasmid #138272, which was digested with SmaI and NotI, using NEB HiFi DNA Assembly Master Mix.

The adenine base editor plasmid (pCMV_ABEmax_P2A_GFP; Addgene #112101), the dCas12-VPR plasmid (Addgene #183956), and the TRE3G-GFP reporter plasmid (Addgene #89453) were obtained from Addgene.

To generate stable cell lines co-expressing ABEmax and T7 RNA polymerase, two PiggyBac (PB) transposon plasmids were constructed. (1) ABE PB plasmid: The ABEmax_P2A_GFP coding sequence was PCR-amplified from pCMV_ABEmax_P2A_GFP (Addgene #112101) and cloned into a modified PB-TRE-dCas9-VPR backbone (Addgene #63800), in which the hygromycin resistance cassette was replaced with puromycin. (2) T7 RNAP PB plasmid: The T7 RNA polymerase coding sequence (lacking an NLS) was PCR-amplified from the cloned T7 plasmid described above and inserted into PB-TRE-Cas9 (Addgene #126029) by replacing the Cas9 coding sequence. The resulting construct retained the hygromycin resistance cassette. All assemblies were performed using Gibson assembly.

### Tissue culture

HEK293T and HeLa cell lines were sourced from the American Type Culture Collection (ATCC, Manassas, Virginia) and maintained at 37 °C and 5% CO_2_. All cell lines were cultured in Gibco™ DMEM with high glucose and pyruvate (ThermoFisher cat. #11-995-073), supplemented with 10% fetal bovine serum (FBS, ThermoFisher cat. #A5209501) and 1% Penicillin-Streptomycin (ThermoFisher catalog #15140122).

### Transient transfection

For the evaluation of different RNAPs with GFP reporter system, approximately 4.5 × 10^4^ HEK293T cells were seeded per well in 48-well tissue culture (TC)-treated plates and cultured to ~80% confluency prior to transfection. Cells were transfected with 100 ng of GFP reporter plasmid and 150 ng of the cognate polymerase plasmid. Transfections were performed using Lipofectamine 3000 (Invitrogen, L3000015). For each well, nucleic acid– Lipofectamine complexes were prepared by combining the desired plasmids with 0.5 µL of Lipofectamine 3000 reagent and 0.5 µL of P3000 enhancer in a total volume of 25 µL Opti-MEM (Thermo Fisher Scientific). Complexes were incubated for 15 minutes at room temperature before being added to the cells.

For evaluating RNAP-dependent gRNA expression from ssDNA templates in the ABE– mCherry reporter system, approximately 7 × 10^4^ HEK293T cells were seeded per well in 24-well, TC–treated plates and grown to ~80% confluency prior to transfection. Cells were transfected with 160 ng of the indicated RNAP plasmid, 200 ng of ABE, 90 ng of the inactive mCherry reporter construct, and 0.5 nM of the corresponding ssDNA template using Lipofectamine 3000 (Thermo Fisher Scientific). For each well, transfection complexes were prepared by combining the plasmids with 1.5 µL Lipofectamine 3000 reagent and 1 µL P3000 enhancer in a total volume of 50 µL Opti-MEM. Complexes were incubated for 15 minutes at room temperature before being added dropwise to the cells.

For assessing ssDNA-driven gRNA expression in HEK293T cell lines stably expressing doxycycline-inducible ABEmax and T7 RNA polymerase, approximately 8 × 10^4^ ABEmax/T7 cells were seeded per well in 24-well, TC–treated plates in media supplemented with 1.5 µg/mL doxycycline (Sigma-Aldrich) and grown to ~80% confluency prior to transfection. Cells were then transfected with 300 ng of the inactive mCherry reporter plasmid and 1 nM ssDNA template using Lipofectamine 3000 (Thermo Fisher Scientific), as described above.

For endogenous genome-editing experiments, approximately 6 × 10^4^ HEK293T cells were seeded per well in 24-well, TC–treated plates and grown to ~70% confluency prior to transfection. Cells were transfected with 200 ng of T7 (no NLS) or Phi15 RNA polymerase plasmid, 250 ng of ABEmax plasmid, and 0.5 nM of the corresponding ssDNA template using Lipofectamine 3000 (Thermo Fisher Scientific), as described above.

For transcriptional activation experiments with dCas12-mVPR in HEK293T cells, 7 × 10^4^ HEK293T cells were seeded per well in 24-well, TC–treated plates and grown to ~80% confluency prior to transfection. Cells were transfected with 160 ng of T7 RNA polymerase plasmid, 200 ng of dCas12-mVPR, 100 ng of the TRE3G-GFP reporter construct, and 1 nM of the corresponding ssDNA template using Lipofectamine 3000 (Thermo Fisher Scientific). For each well, transfection complexes were prepared by combining the plasmids with 1.5 µL Lipofectamine 3000 reagent and 1 µL P3000 enhancer in a total volume of 50 µL Opti-MEM. Complexes were incubated for 15 minutes at room temperature before being added dropwise to the cells.

For transcriptional activation experiments with dCas12-mVPR in HeLa cells, 7 × 10^4^ HeLa cells were seeded per well in 24-well, TC–treated plates and grown to ~80% confluency prior to transfection. Cells were transfected with 80 ng of T7 RNA polymerase plasmid, 100 ng of dCas12-mVPR, 50 ng of the TRE3G-GFP reporter construct, and 0.3 nM of the corresponding ssDNA template using Lipofectamine 3000 (Thermo Fisher Scientific). For each well, transfection complexes were prepared by combining the plasmids with 0.75 µL Lipofectamine 3000 reagent and 1 µL P3000 enhancer in a total volume of 50 µL Opti-MEM. Complexes were incubated for 15 minutes at room temperature before being added dropwise to the cells.

### Cell line generation

HEK293T cells were seeded in 6-well TC-treated plates and cultured until approximately 75% confluency before transfection. Cells were transfected with 2.5 µg of the cloned piggyBac plasmids (as described above) and 1 µg of Super piggyBac Transposase plasmid (System Biosciences, PB210PA-1) using Lipofectamine 3000. 48 hours post-transfection, selection was initiated by adding 2 µg/mL puromycin and 25 µg/mL hygromycin to the culture medium. Cells were maintained under these selective conditions for two weeks to ensure stable integration and selection of transfected cells.

### Genomic DNA processing and sequencing

Genomic DNA was extracted 60 hours after transfection using a Zymo Research Quick-DNA Microprep Kit (Zymo Research, D3020). Genomic areas of interest were amplified from 6 ng of whole genome DNA using Q5^®^ Hot Start High-Fidelity 2X Master Mix (NEB, 0494) according to the manufacturer’s protocol. This was followed by DNA separation via electrophoresis on 1% E-Gel EX Agarose Gels (Invitrogen) and extraction using a Monarch DNA Gel Extraction Kit (NEB, T1020S). Sanger sequencing was performed using Azenta Genewiz.

### Fluorescence Microscopy

Fluorescent signals within the cells were detected and analyzed using the FLoid™ Cell Imaging Station (Life Technologies).

### Flow cytometry analysis

48 or 72 hours post transfection, cells were harvested by removing the media, washed one time with PBS, and then trypsinized using TrypLE Express (Thermo Fisher Scientific, 12604013) for 3 minutes. Resuspended cells were pelleted via centrifugation for 5 minutes at 500 g. After centrifugation, cell pellets were resuspended into PBS with 2% FBS and transferred into 5 ml round bottom polystyrene flow tubes with cell strainer (Corning Life Sciences, 352235). Cells were analyzed by flow cytometry (Symphony A1, BD Biosciences) and data was processed with FlowJo^*TM*^ 10 software.

## Acknowledgments

G.M.C., A.M., and R.F. were supported by the Aging and Longevity-Related Research Fund. A.M. was supported by the Ibn Rushd Postdoctoral Fellowship from King Abdullah University of Science and Technology.

## Author Contributions

A.M. Conceptualized and initiated the study. A.M., R.M.P., R.O., and R.F. performed experiments. A.M., R.M.P., and R.F. analyzed the data. A.M. and G.M.C. supervised the project. A.M. and R.F. wrote the manuscript with input from all authors.

## Competing Interests

The authors declare no competing interests. Full disclosures for G.M.C. can be found at https://arep.med.harvard.edu/t/

## Notes

### Competing Interest Statement

The authors have declared no competing interest.

